# Molecular diagnostics for early detection of invasion of malaria vector *Anopheles stephensi*

**DOI:** 10.1101/2022.09.26.509490

**Authors:** Om P. Singh, Taranjeet Kaur, Gunjan Sharma, Kona M. Prasad, Neera Kapoor, Prashant K. Mallick

## Abstract

There are increasing reports of the expansion of the Asian malaria vector *Anopheles stephensi* in new geographical areas posing a threat to the elimination of urban malaria. Efficient surveillance of this vector in affected areas and early detection in new geographical areas is key for the containment and control of this species. To overcome the practical difficulties associated with the morphological identification of immature stages and adults of *An. stephensi*, a species-specific PCR and a Real-Time PCR assay were developed for their identification targeting a unique segment of ITS2 lacking homology to any other organism. Both PCRs can be used to identify *An. stephensi* individually or in pooled samples of mixed species, including when present in extremely low proportion. This study also reports a method for selective amplification and sequencing of *An. stephensi* for their confirmation in pooled samples of mixed species.

## Introduction

There are increasing reports of the expansion of the Asian malaria vector *Anopheles stephensi* [1-6] with an increased threat of urban malaria [7]. The gradual southward expansion of this species was recorded in India since the 1970s [1]. The first-ever occurrence of *An. stephensi* in the African subcontinent was reported as early as 1966 in a town in Egypt [2]. During the last two decades, several reports of expansion of this species in the countries of the Horn of Africa, the Republic of Sudan, Sri Lanka and Lakshadweep Islands (Indian union territory) have appeared [1-6]. A high probability of presence within many urban cities across Africa has been predicted, which warranted the prioritization of vector surveillance [8]. As a consequence of recent invasions of this vector species in several countries, World Health Organization recommended conducting active surveillance of *An. stephensi* in urban and peri-urban areas, in addition to routine surveillance in rural areas in the affected and surrounding geographical areas [9]. However, the identification of this species, which relies mainly on the morphological characters of adult-female mosquitoes, is often challenging. The sampling of adult *An. stephensi* from their resting habitats is difficult because it is secretive in its habits [10]. World Health Organization (WHO) has recommended sampling of immatures from natural breeding habitats and rearing them in the laboratory until their emergence into adults to facilitate species identification [9]. Such practice is being adopted for sampling this species in African regions [4, 5, 11], which is a time-consuming and labour-intensive process. Even freshly emerged adults have been reported to be misidentified as *An. gambiae* due to superficial resemblance [11]. Adult mosquito collection through light-trap or pyrethrum spray collections (PSC) are alternative and popular methods of sampling *An. stephensi* but the identification of adult mosquitoes collected through such methods may be difficult due to the loss of morphological characteristics critical for their correct identification. Therefore, the development of highly specific PCR-based assays is crucial for the identification of both larval and adult *An. stephensi* species collected by a variety of methods. Such diagnostics will be very helpful to field technologists who are not familiar with the morphology of this species. These assays can detect *An. stephensi* in a large pool of mosquitoes (e.g., 1 in 500) when their proportion is extremely low. Equally important is the development of a DNA sequencing strategy to confirm the presence of *An. stephensi* in such pools of mixed species. These molecular tools will serve as an important tool for the early detection of invasions of *An. stephensi* in a new geographical area where it is present in extremely low densities.

## Methods

### Mosquito samples

The list of mosquitoes and preserved DNA samples used in this study is provided in **Table 1**. The field-collected *An. stephensi* were processed for the identification of biological forms following Singh et al. [12].

**Table 1.**
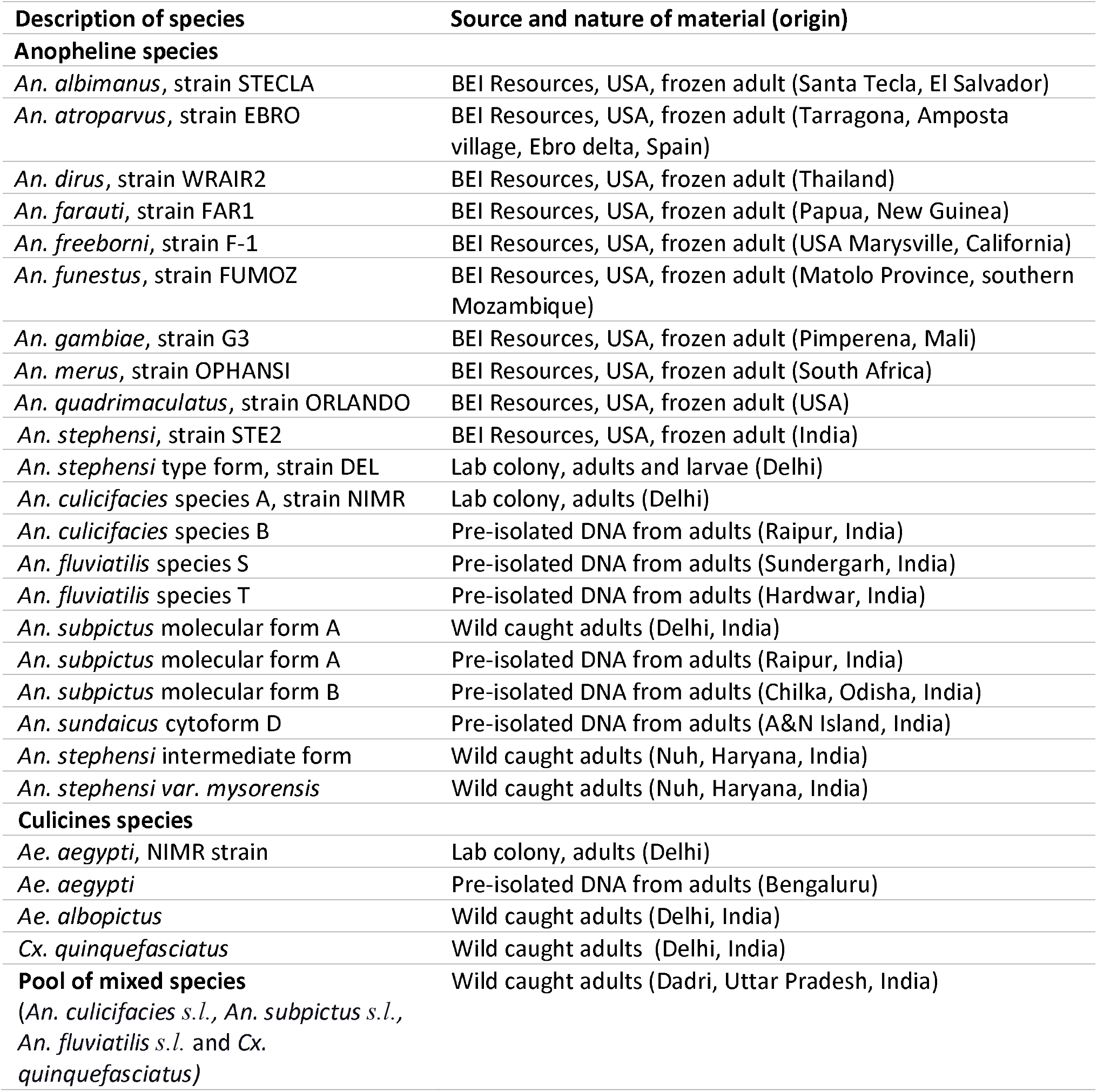
List of biological material used in the study.

### DNA isolation from individual and pooled samples

DNA was isolated from the individual as well as pooled samples of mosquitoes for standardization of PCR-based assays and their validation. Pooled samples of different sizes were used, each comprising a single *An. stephensi* and rest *An. culicifacies*. A pool of field-collected mosquitoes comprising single *An. stephensi* and rest other species was also used.

For DNA isolation, two commercial kits were selected. For DNA isolation from individual mosquitoes or smaller pool sizes, DNeasy Blood & Tissue kit (Qiagen Inc), recommended for ≤25 mg of tissue, was used. For larger pools, DNAzol Reagent (Thermo Fisher Scientific), recommended for 25-50 mg of tissue per ml of reagent, was used. In addition, DNA was isolated from individual *An. stephensi* by boiling method.

#### 1. DNeasy Blood & Tissue Kits

DNA was isolated from individual mosquitoes of some Anopheles, Culex and Aedes species (listed in **Table 1**.) following manufacturer’s protocol and eluted in 200 μL elution buffer. DNA from smaller pools of III-IV instar larvae or adults, each containing a single *An. stephensi* and the rest *An. culicifacies* in different pool sizes, i.e., 2, 5, 10, 20 and 50, were isolated. DNA from single leg of *An. stephensi* mosquitoes was also isolated using this kit which was eluted in 50 μL of elution buffer.

#### 2. DNAzol Reagent

DNAzol Reagent was used for DNA isolation from pools of adult (i.e., 25, 100 and 500) mosquitoes and pools of 100 larvae (III-IV instar), each containing one *An. stephensi* and the rest *An. culicifacies*. Pools of 25 and 100 mosquitoes were directly homogenized in 1 ml of DNAzol reagent in a microcentrifuge tube. In the case of pools of 100 mosquitoes, 250 μL of the triturate was transferred to a separate microcentrifuge tube and made up a volume of 1 ml with DNAzol. The pools of 500 mosquitoes were ground in liquid nitrogen and approximately 25 mg of triturate was transferred in a microcentrifuge tube and homogenized in 1 ml of DNAzol reagent. All triturates were centrifuged at 10,000 × g for 10 min and 500 μl of supernatant was transferred to a fresh 1.5 ml microcentrifuge tube which was subjected to ethanol precipitation, washing and solubilization of DNA following the vendor’s protocol. DNA was dissolved in 200 μL of 8.0 mM NaOH. DNA was also isolated from a pool of 100 mosquitoes containing a single *An. stephensi* and rest field-collected (through hand-catch method) mosquitoes belonging to *An. culicifacies, An. subpictus, An. fluviatilis* and *Cx. quinquefasciatus*.

#### 3. Boiling method

DNA from 10 individual specimens of *An. stephensi* were also isolated by boiling method following Sharma et al. [13].

### Selection of target sites for designing primers and probes

Ribosomal DNA (rDNA) was selected as a target site for developing diagnostics for the identification of *An. stephensi*, which are present in hundreds of copies in an individual and are highly conserved in a species due to homogenization of sequence through unequal crossing over and gene conversion, a process known as ‘concerted evolution’. The second internal transcribed spacer (ITS2)-rDNA, which is conserved within a species but highly variable across taxa, was selected for designing *An. stephensi*-specific primers and probes. For designing *An. stephensi*-specific primers and probes, a homology search of ITS2 sequences of *An. stephensi* [14] was done through NCBI’s nucleotide BLAST-search utility. The search was optimized for blastn (somewhat similar sequences) and the taxon ‘*Anopheles stephensi*’ was excluded from the search. All 297 search returns belonged to the *Anopheles* mosquitoes, all belonging to the *Neocellia* series (subgenus *Cellia*); however, none of the returns showed homology to the last 122 bp segment of ITS2 (**Figure 1**). We considered this region unique to *An. stephensi* and exploited this region for designing highly-specific *An. stephensi*-specific primers and probes. For designing universal primers and a probe, highly conserved regions were identified based on the alignment of 5.8S and 28S rDNA sequences of anophelines available in the GenBank. The locations of primers and PCR strategies are displayed in **Figure 1** and the list of primers and probes is shown in **Table 2**. The specificity of each *An. stephensi*-specific primer and probes were checked in silico by performing NCBI-BLAST search utility (blastn) and ensured that none of these matched with rDNA sequence of any other mosquitoes.

**Table 2.**
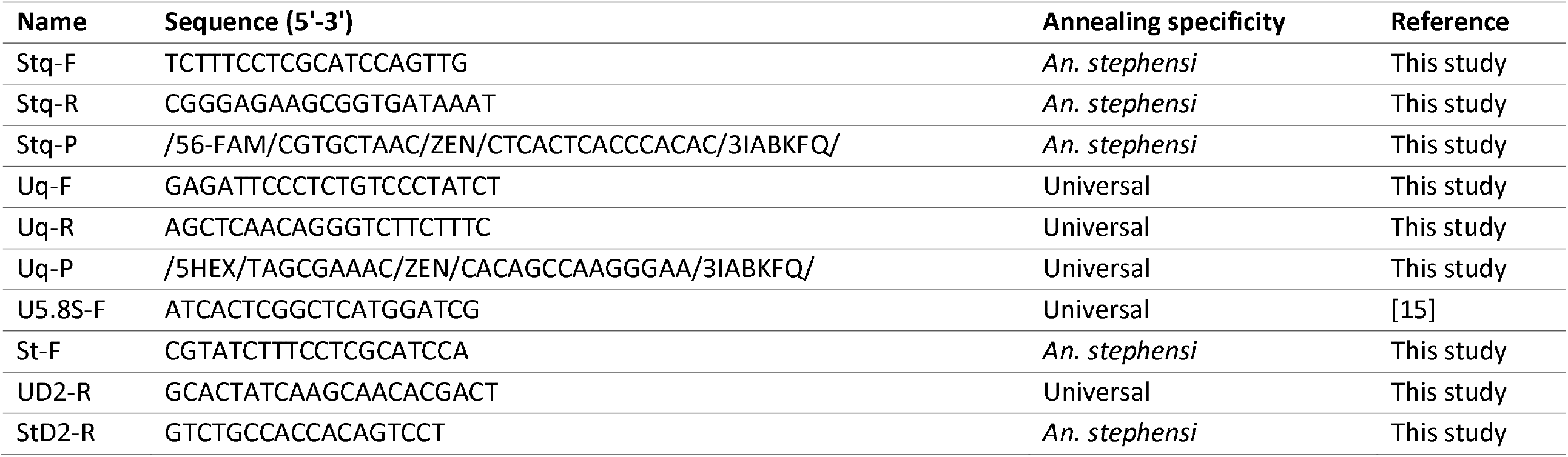
Sequences of primers and probes used.

**Figure 1.**
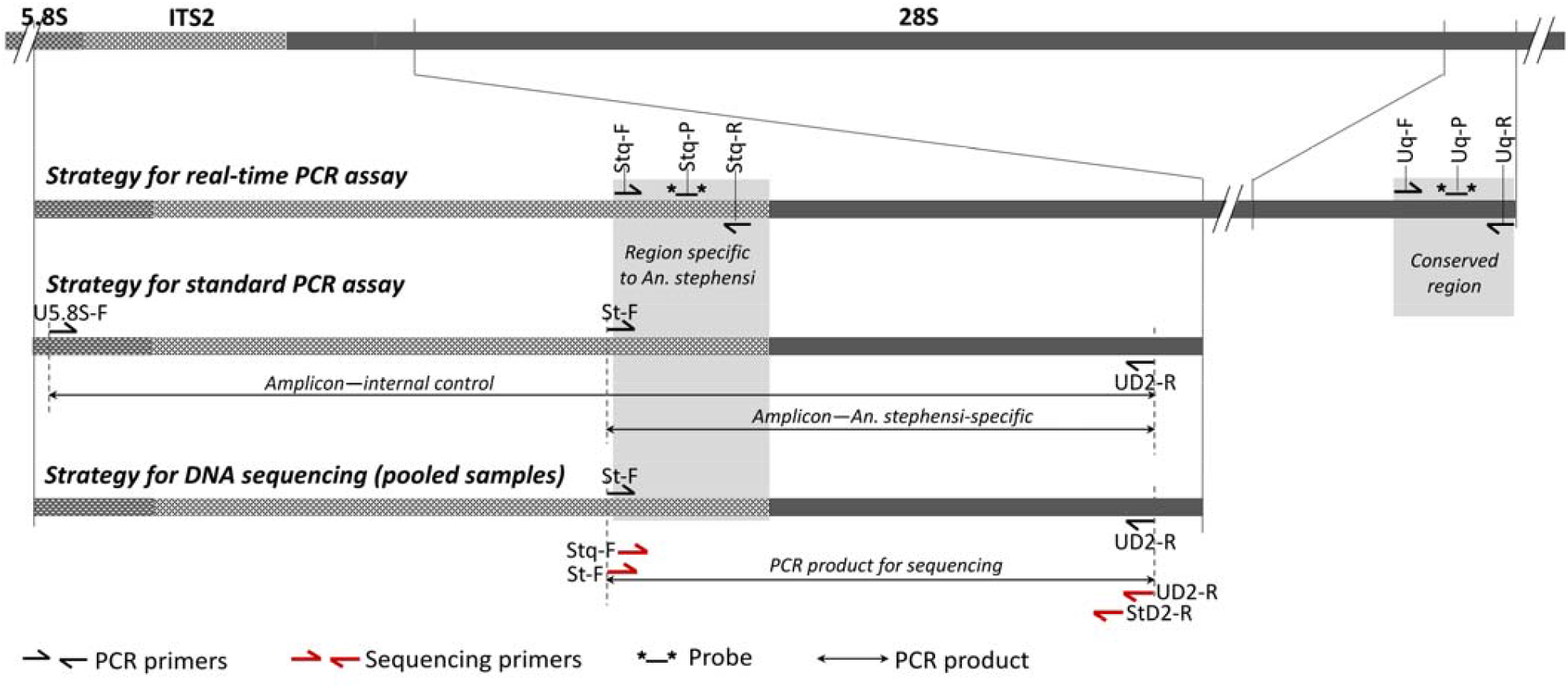
Schematic representation of PCR strategies

### Development of a hydrolysis real-time PCR assay

For *An. stephensi*-specific hydrolysis real-time PCR assay, two oligonucleotide primers, Stq-F and Stq-R, and a hydrolysis probe Stq-P were designed from the stephensi-specific ITS2 region. For internal control (IC), primers, Uq-F and Uq-R, and a hydrolysis probe Uq-P, were designed from a region of 28S-rDNA conserved in anophelines. The sequences of primers/probes have been provided in **Table 2**. Real-time PCR assays were performed in 10 μL of reaction mixture containing 0.4 μM of each of Uq-F, Uq-R and Stq-F; 0.5 μM of Stq-R; 0.2 μM of each probe (Stq-P and Uq-P); 1X QuantiFast Multiplex PCR kit (Qiagen Inc) and 1 μL of template DNA in Bio-Rad CFX96 Touch Real-Time PCR Detection System. The cyclic conditions were: pre-denaturation at 95°C for 5 mins, followed by 35 cycles, each with denaturation at 95°C for 10 sec and annealing/extension at 60°C for 30 seconds. The number of cycles required for the fluorescent signal to cross the threshold (Ct) values were scored by the software CFX Maestro (**Figure 2**).

**Figure 2.**
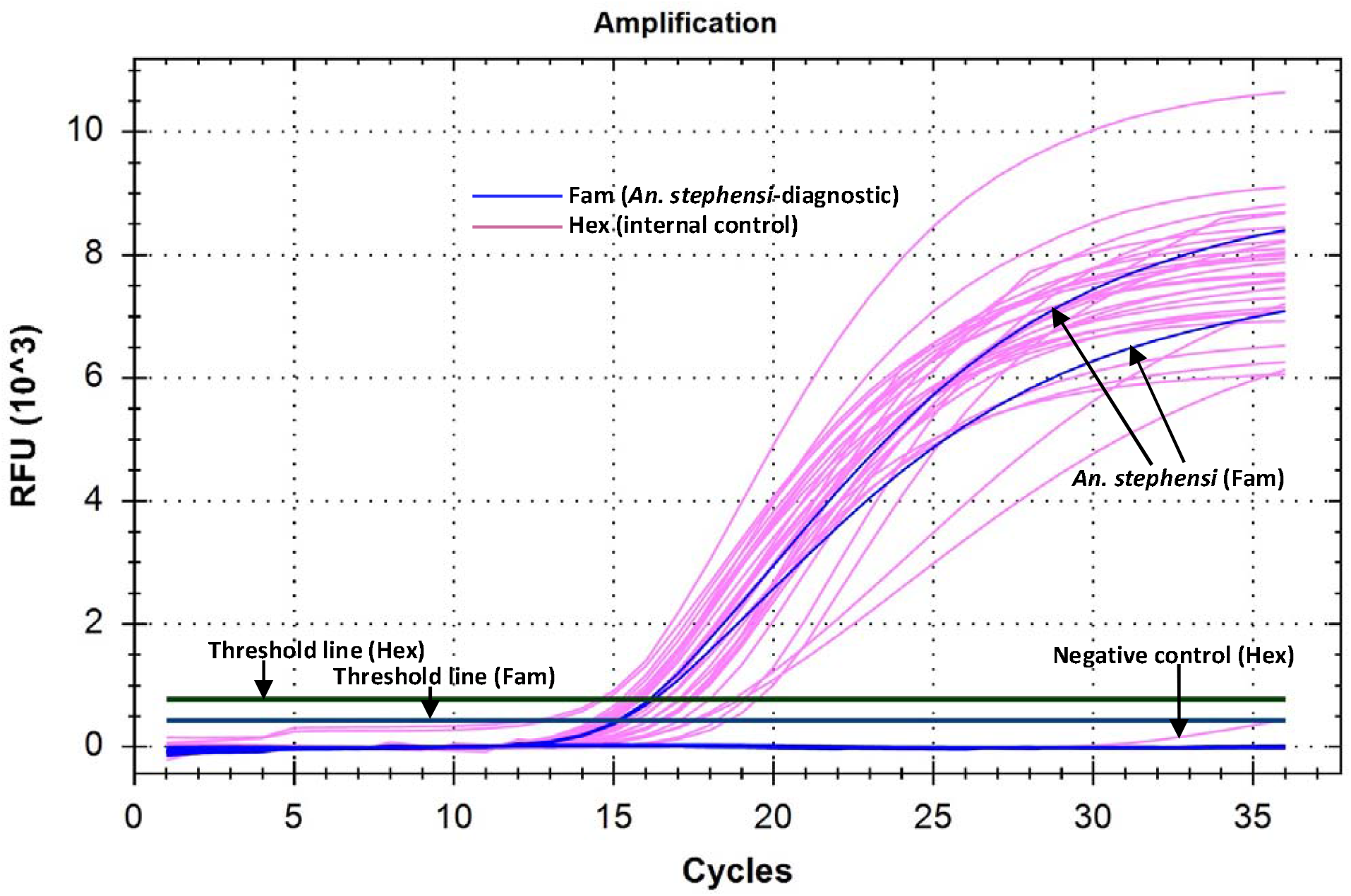
Amplification curve of real-time PCR on *An. stephensi* and other mosquitoes (anophelines and culicine) showing threshold lines for Fam and Hex as determine by the software. Amplification crossing threshold value for Fam was seen in *An. stephensi* only and for Hex was seen in all mosquitoes tested (*An. stephensi, An. albimanus, An. quadrimaculatus, An. dirus, An. farauti, An. gambiae, An. freeborni, An. funestus, An. atroparvus, An. culicifacies* complex, *An. subpictus* complex, *An. merus, An. fluviatilis* complex, *Cx. quinquefasciatus*).

PCR efficiencies of each hydrolysis probe assay were evaluated by performing duplex real-time PCR assays in triplicate at six different concentrations, diluted serially by 10-fold. The experiments were performed on two samples of *An. stephensi*-DNA with different DNA concentrations, 8.7 and 3.2 ng/μL (Sample A and B, respectively in **Figure 3**). To detect the sensitivity (limit of detection), real-time PCR was carried out on the diluted DNA of *An. stephensi* with concentrations of 160 fg, 80 fg, 40 fg, and 20 fg, each with 12 replicates.

**Figure 3.**
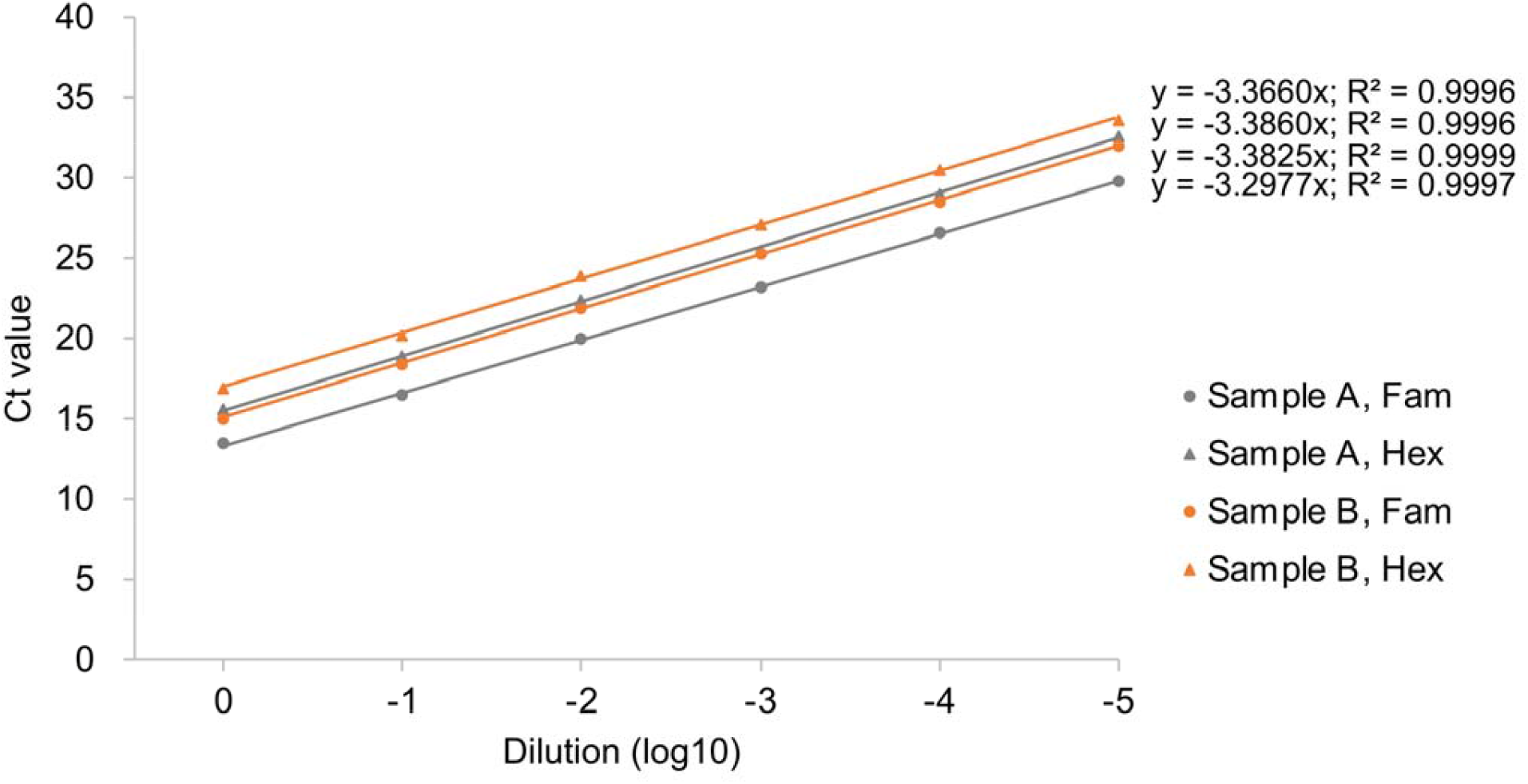
Standard curve showing correlation of number of cycles required for the fluorescent signal to cross the threshold (Ct) values against 10-fold serially diluted DNA samples of *An. stephensi* (two samples, A and B) in the duplex hydrolysis fluorescent probe assay. The slope of each line represents [− 1/log_10_ (PCR efficiency)] for a hydrolysis probe assay. R^2^ represents correlation coefficient of a slope.

### Development of size-diagnostic PCR assay

For a size-diagnostic PCR assay, three primers were designed: the *An. stephensi*-specific forward primer, St-F, was designed from the *An. stephensi*-specific region of ITS2, and two flanking universal primers, U5.8S-F and UD2-R, were designed from conserved 5.8S rDNA and D2 domain of 28S rDNA, respectively. According to the strategy designed (**Figure 1**), a stephensi-diagnostic amplicon of 438 bp size will be formed by the primers St-F and D2-R; and a universal amplicon of varying sizes (>600 bp), depending upon the length of ITS2 in a particular species, will be formed by the primers 5.8S and D2-R. The universal amplicon will serve as an IC to rule out PCR failure.

Due to the competitive nature of primers in multiplex-PCR, two protocols of PCR were optimized based on the number of mosquitoes in a pool. For individual mosquitoes or smaller pools, end-point PCR assays were carried out with Hot Start Taq 2X Master Mix (New England Biolabs, USA) in a 20 μL reaction mixture containing 0.5 units of taq polymerase, 1.5 mM of MgCl_2_, 0.25 μM of primer St-F, 0.25 μM of primer U5.8S-F, 0.375 μM of UD2-R, and 0.50 μL of DNA template (Protocol-1). In another protocol (Protocol-2), the concentration of primer U5.8S-F was reduced to 0.10 μM and concentration of UD2-R was increased to 0.50 μM for larger pools of mosquitoes (25-500). It was also observed that the intensity of amplicon in size-diagnostic PCR with pools of mosquitoes (>25) could be improved by the dilution of template DNA. Therefore, the template DNA of larger pools of mosquitoes (>25) was further diluted by 1/10 before using it as template DNA for size-diagnostic PCR reactions in order to minimize the PCR inhibitors in PCR reactions. However, such dilution was not required for real-time PCR. Optimized thermal cycling conditions were: an initial denaturation step at 95 °C for 30 sec, 30 cycles, each with a denaturation step at 95 °C for 30 sec, annealing at 55 °C for 30 sec and extension at 68 °C for 45 sec, and final extension at 68 °C for 7 min. Five μL of PCR product was run on 2% agarose gel and visualized in the gel documentation system.

### DNA sequencing strategy for the confirmation of PCR-based identification of *An. stephensi* in pooled samples

*Anopheles stephensi* targeted amplicons were amplified from DNA isolated from pools of 100 (field-collected) and 500 mosquitoes, each pool containing a single *An. stephensi*, using primers St-F and UD2-R. PCR was performed using Hot Start Taq 2X Master Mix (New England Biolabs, USA) in a 20 μL reaction mixture containing 0.25 μM of each primer. PCR conditions were similar to PCR protocol-1 except the extension time, which was reduced to 30 sec and number of cycles was increased to 35. The amplified products were treated with Exo-Sap II (Thermo Fisher Scientific), and sequence termination reactions were performed from both directions of strands using BigDye Terminator v3.2 (Applied Biosystem). The primers used for sequencing were the primers used for PCR amplification as well as the two internal primers (Stq-F and StD2-R, **Table 2**). Both internal primers are specific to *An. stephensi* and expected to provide a noise-free sequence by eliminating the possibility of sequencing non-specific PCR product. The sequencing products were electrophoresed on a ABI Prism 3730xl.

## Results

### Real-time PCR assay

PCR efficiencies of *An. stephensi* diagnostic (Fam-labelled probe) and IC (Hex-labelled probe) PCRs as estimated based on Ct values of six serially diluted concentrations of DNA were 97.5-101% and 97.4-98.2%, respectively. The slopes and correlation coefficients is displayed in **Figure 2**. The dynamic range of Ct values for real-time PCR for *An. stephensi*-specific PCR was 13.5 to 32 and for IC-PCR was 15.5 to 33.5 and the limit of detection (LOD) was 40 fg of genomic DNA.

The results of the real-time PCR assays carried out on DNA isolated from individual *An. stephensi* samples, 19 different non-target Anopheles species and a single *An. stephensi* in different pool sizes are provided in **Table 3**. Both hydrolysis real-time assays were specific based on software determined Ct value scored within 35 cycles of reactions, except in case of two pre-isolated DNA (one each of Ae. aegypti and *An. subpictus*) showing false positivity with late Ct values (>32). These two samples were contaminated with the *An. stephensi* DNA as revealed through *An. stephensi*-targeted sequencing. The sequencing methodology has been provided in **Supplementary File S1**). Although primers/probe for IC was designed based on anophelines’ 28S rDNA sequences, they also worked on all three non-anopheline species tested, i.e., *Culex quinquefasciatus, Aedes aegypti* and *Ae. albopictus*. The real-time PCR also successfully identified *An. stephensi* from DNA samples isolated by boiling method and DNA isolated from a single leg of mosquitoes. The real-time PCR was sensitive to detecting single *An. stephensi* in pools of 500 mosquitoes with low Ct values (20.12 and 23.96). Rising of florescent signal was noticed with internal control probe in some experiments only in negative controls after 30 cycles but not with stephensi-specific probe (**Figure 2**).

**Table 3.** Results of hydrolysis real-time PCR and size-diagnostic PCR assays on individual and pooled mosquitoes.

|  | DNA isolation method | Real-Time PCR-assay<sup>§</sup> |  |  | Size-diagnostic PCR-assay |  |  |
| --- | --- | --- | --- | --- | --- | --- | --- |
|  |  | n | Ct values (*An. stephensi*) | Ct values (IC) | n | *An. stephensi*-specific band | IC band |
| **A. Anophelines** |  |  |  |  |  |  |  |
| ***Neocellia* Series** |  |  |  |  |  |  |  |
| *An. stephensi* (type) | DNeasy | 26 | 13.47-17.23 | 15.20-18.93 | 24 | Positive | Positive |
| *An. stephensi* (intermediate) | DNeasy | 2 | 14.75-15.60 | 15.39-16.06 | 2 | Positive | Positive |
| *An. stephensi* (var. *mysorensis*) | DNeasy | 1 | 14.54 | 15.01 | 1 | Positive | Positive |
| *An. stephensi* Strain STE2 | DNeasy | 1 | 14.85-15.25 | 16.06-16.51 | 2 | Positive | Positive |
| *An. stephensi* (type form) | Boiling | 10 | 15.17-17.41 | 15.97-18.05 | - | Not done | Not done |
| *An. stephensi* (type form, single leg) | DNeasy | 4 | 18.64-22.29 | 19.02-23.51 | 4 | Positive | positive |
| ***Pyretophorus* Series** |  |  |  |  |  |  |  |
| *An. gambiae* | DNeasy | 3 | Negative | 14.58-17.32 | 3 | Negative | positive |
| *An. quadrimaculatus* | DNeasy | 2 | Negative | 14.52-15.64 | 2 | Negative | positive |
| *An. merus* | DNeasy | 2 | Negative | 15.22-15.35 | 2 | Negative | positive |
| *An. subpictus* form A | DNeasy | 2 | Negative | 16.61-17.02 | 1 | Negative | positive |
| *An. subpictus* form A<sup>¶</sup> | Pre-isolated<sup>†</sup> | 1 | 33.09 | 18.32 |  | Negative | positive |
| *An. subpictus* form B | Pre-isolated<sup>†</sup> | 2 | Negative | 16.17-16.35 | 1 | Negative | positive |
| *An. sundaicus* cytoform D | Pre-isolated<sup>†</sup> | 1 | Negative | 14.47 | 1 | Negative | positive |
| ***Albimanus* Series** |  |  |  |  |  |  |  |
| *An. albimanus* | DNeasy | 2 | Negative | 16.21-16.58 | 1 | Negative | Positive |
| ***Anopheles* Series** |  |  |  |  |  |  |  |
| *An. freeborni* | DNeasy | 2 | Negative | 15.73-15.81 | 1 | Negative | Positive |
| *An. atroparvus* | DNeasy | 2 | Negative | 14.70-15.88 | 1 | Negative | Positive |
| ***Neomyzomyia* Series** |  |  |  |  |  |  |  |
| *An. dirus* | DNeasy | 2 | Negative | 17.34-18.01 | 1 | Negative | Positive |
| *An. farauti* | DNeasy | 2 | Negative | 19.09-19.48 | 1 | Negative | Positive |
| ***Myzomyia* Series** |  |  |  |  |  |  |  |
| *An. funestus* | DNeasy | 2 | Negative | 15.50-15.62 | 1 | Negative | Positive |
| *An. culicifacies* species A | DNeasy | 1 | Negative | 17.41 | 1 | Negative | Positive |
| *An. culicifacies* species B | Pre-isolated<sup>†</sup> | 1 | Negative | 17.48 | 1 | Negative | Positive |
| *An. fluviatilis* species S | Pre-isolated† | 1 | Negative | 16.61 | 1 | Negative | Positive |
| *An. fluviatilis* species T | Pre-isolated† | 1 | Negative | 17.02 | 1 | Negative | Positive |
| **B. Culicines** |  |  |  |  |  |  |  |
| *Cx. quinquefasciatus* | DNeasy | 3 | Negative | 16.58-19.20 | 2 | Negative | Positive |
| *Ae. albopictus* | DNeasy | 2 | Negative | 16.65-16.68 | 2 | Negative | Positive |
| *Ae. aegypti* | DNeasy | 2 | Negative | 18.96-19.20 | 2 | Negative | Positive |
| *Ae. aegypti*<sup>¶</sup> | Pre-isolated† | 1 | 32.95 | 20.97 |  | Negative | positive |
| **C. Single *An. stephensi* individual pooled with *An. culicifacies*\*** |  |  |  |  |  |  |  |
| Adults (1/2) | DNeasy | 1 | 17.06 | 18.31 | 1 | Positive | Positive |
| Adults (1/5) | DNeasy | 1 | 15.37 | 16.11 | 1 | Positive | Positive |
| Adults (1/10) | DNeasy | 1 | 15.77 | 15.30 | 1 | Positive | Positive |
| Adults (1/20) | DNeasy | 1 | 15.74 | 13.71 | 1 | Positive | Positive |
| Adults (1/50) | DNeasy | 1 | 15.43 | 11.43 | 1 | Positive | Positive |
| Adults (1/25) | DNAzol | 1 | 14.53 | 12.61 | 1 | Positive | Positive |
| Adults (1/100) | DNAzol | 1 | 16.37 | 11.27 | 1 | Positive | Positive |
| Adults (1/500) | DNAzol | 2 | 20.12-23.96 | 11.08-11.50 | 2 | Positive | Positive |
| Larvae (1/2) | DNeasy | 1 | 15.56 | 16.87 | 1 | Positive | Positive |
| Larvae (1/5) | DNeasy | 1 | 15.49 | 16.65 | 1 | Positive | Positive |
| Larvae (1/10) | DNeasy | 1 | 16.36 | 15.53 | 1 | Positive | Positive |
| Larvae (1/20) | DNeasy | 1 | 16.51 | 15.06 | 1 | Positive | Positive |
| Larvae (1/50) | DNeasy | 1 | 16.27 | 14.40 | 1 | Positive | Positive |
| Larvae (1/100) | DNAzol | 1 | 18.14 | 13.16 | 1 | Positive | Positive |
| **D. Single *An. stephensi* individual pooled with field collected mosquitoes (mixed mosquito-species)\*** |  |  |  |  |  |  |  |
| Adults (1/100) | DNAzol | 1 | 17.61 | 11.70 | 1 | Positive | Positive |

<sup>§</sup>Ct value shown as “Negative” implies that no Ct value was scored within 35 cycles of reactions. \*Denominator of numeric expression inside parenthesis indicate total number of mosquitoes in a pool. †Preserved DNA isolated manually for some other studies. <sup>¶</sup>Contaminated with *An. stephensi*-DNA

### Size-diagnostic PCR assay

PCR protocol-1 was performed on DNA samples isolated from individual *An. stephensi* samples, non-target mosquitoes and pooled samples of mixed species (up to 50 mosquitoes) which provided desired amplicons. All the *An. stephensi* were positive for *An. stephensi*-specific band (438 bp) and all other 19 mosquito species were negative (**Table 3; Figure 4 & Figure 5**). All species exhibited amplification of an IC band of varying sizes (>600 bp). *Ae. aegypti* exhibited the smallest IC band (∼650 bp) due to the shortest length of ITS2 (200 bp). *An. funestus* and *An. dirus* exhibited largest IC bands (>1 kb) due to longer ITS2 (700 bp). PCR protocol-1 successfully identified *An. stephensi* in all pooled samples of mixed species, where *An. stephensi*-specific band was prominent in pools of up to 20 mosquitoes. It was observed that the *An. stephensi* diagnostic band gets fainter as the concentration of *An. stephensi* DNA decreases in larger pools (**Figure 4A**). Therefore, a different protocol (Protocol-2) was adopted for larger pools with different primer concentrations. PCR protocol-2 successfully identified *An. stephensi* in pools of 25 to 500 mosquitoes, and provided clearly visible *An. stephensi*-specific band in pools of up to 100 mosquitoes. Based on the results, we found PCR protocol-1 suitable for individual samples or pools of smaller size (up to 25) (**Figure 4A**) and PCR protocol-2 for larger pools (25-100 mosquitoes) (**Figure 4B & C**).

**Figure 4.**
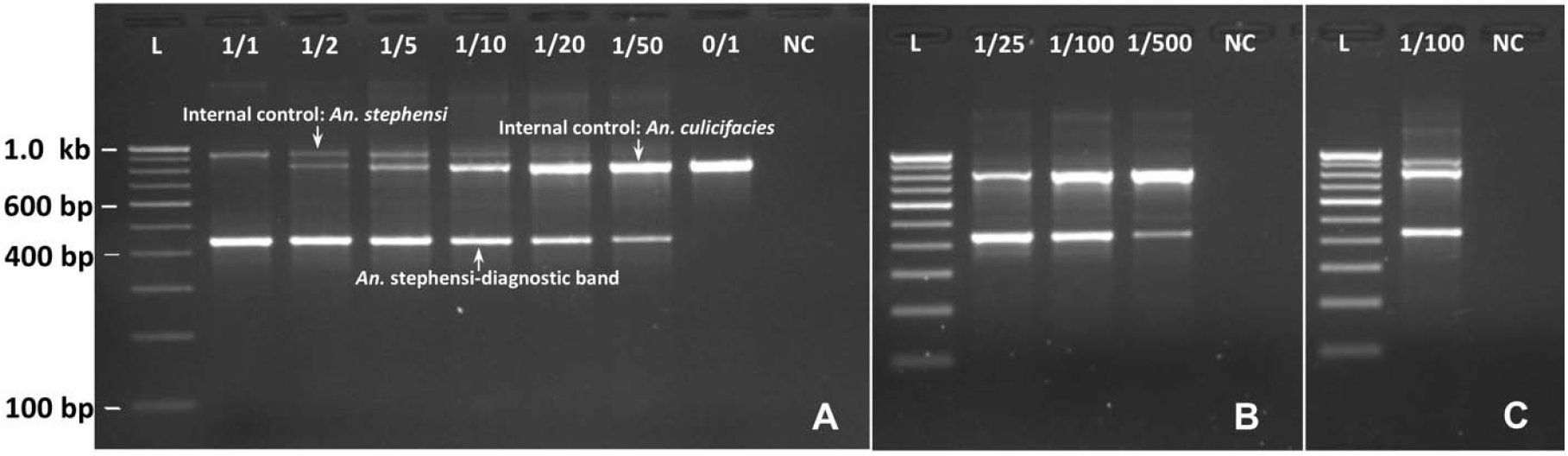
Gel photographs visualizing the result of *An. stephensi*-specific PCRs. A: PCR protocol-1 on individual and pools of *An. stephensi* and *An. culicifacies*. B: PCR protocol-2 on larger pools of *An. stephensi* and *An. culicifacies*. C: PCR protocol-2 on a pool of 100 mosquitoes containing a single *An. stephensi* mixed with other wild-caught anophelines (*An. culicifacies, An. fluviatilis, An. minimus* and *An. subpictus*). The numerator of numeric expression shown on the top of each lane indicates number of *An. stephensi* in a pool, and denominator indicates size of mosquito-pool. L= 100 bp ladder; NC=negative control.

**Figure 5.**
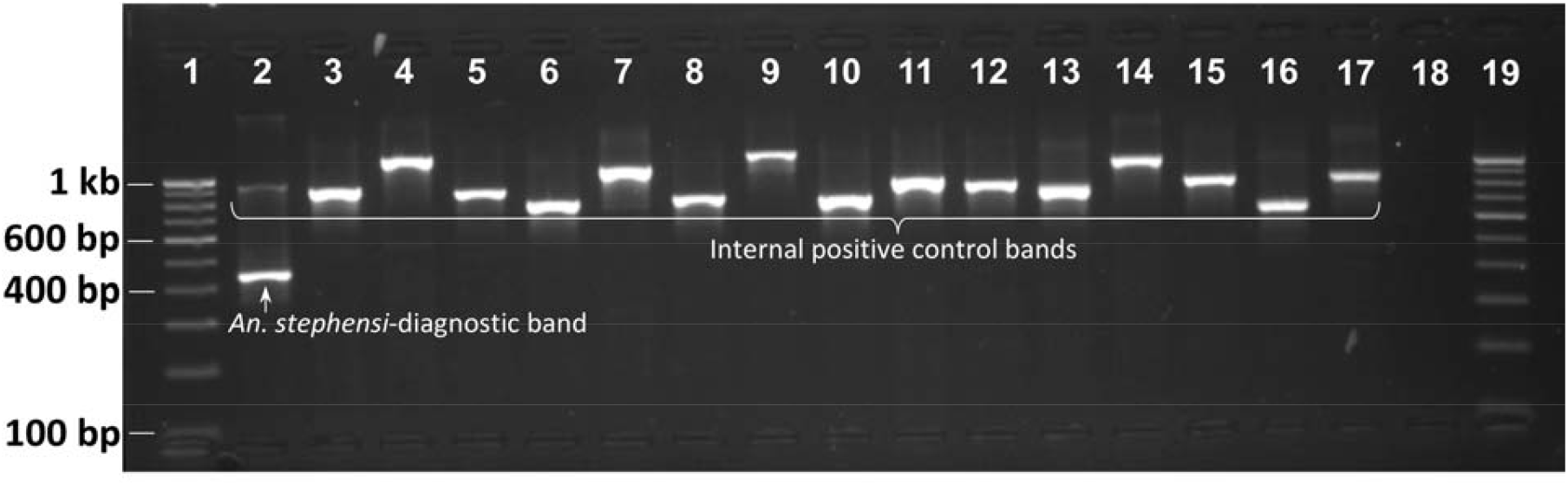
Gel photograph showing the result of *An. stephensi*-specific PCR (protocol-1) on some individual mosquitoes belonging to the genus *Anopheles, Culex* and *Aedes*. Lanes 1 & 19: 100 bp DNA ladder; 2: *An. stephensi*; 3: *An. gambiae*; 4: *An. dirus*; 5: *An. albimanus*; 6: *An. quadrimaculatus*; 7: *An. farauti*; 8: *An. freeborni*; 9: *An. funestus*; 10: *An. atroparvus*; 11: *An. merus*; 12: *An. fluviatilis* species T; 13: *Cx. quinquefasciatus*; 14: *An. subpictus* molecular form A; 15: *An. minimus s.s.*; 16: *Ae. aegypti*; 17: *Ae. albopictus*; 18: negative control.

### Sequencing results

*Anopheles stephensi*-targeted DNA sequencing was successful with all four sequencing primers. The quality of DNA sequences generated from pooled mosquitoes was reasonably high (**Figure S1**). The output sequences showed 100% similarity with *An. stephensi* sequences in a BLAST search. The second highest similarity was with *An. superpictus* with 94.88% similarity based on only 73% coverage. The remaining 27% nucleotide sequence belonging to the ITS2 region did not show a match with any organism other than *An. stephensi*.

## Discussion

The molecular methods developed in this study can identify and confirm *An. stephensi* individually or in a large pool of mixed mosquito species, which may facilitate screening large numbers of samples collected from a variety of methods (such as light trap, PSC, larval collection, etc.) with relatively limited effort and time. This will help in early reporting of the presence of *An. stephensi* to concerned state health agencies and the World Health Organization.

In molecular diagnostics, *An. stephensi*-specific primers and probes were designed from a segment of ITS2 lacking homology to any other organisms for which the rDNA sequence database is available in the public domain, enabling the design of primers that are highly specific to *An. stephensi* and refractory to nonspecific annealing. However, this doesn’t preclude the possibility of any mosquito’s sequence, unreported yet, having matching sequences with *An. stephensi*-specific primers/probe. Therefore, it is suggested that confirmatory DNA sequencing must be performed in new areas using stephensi-specific primers suggested in this report. In earlier studies, confirmation of this species was done through sequencing of ITS2 and mitochondrial DNA using universal primers, which cannot be used in pooled samples of mixed species. Moreover, Mishra et al. [14] have shown that direct DNA sequencing of ITS2 in the case of *An. stephensi* is not fruitful due to the presence of two haplotypes in individuals which differs by 2 bp indel which causes the collapse of a sequence starting from the indel position. The method proposed here for sequencing *An. stephensi* in a pooled sample was targeted to indel-free partial ITS2 (which lacks homology with any organism) and D1-D2 domains of 28S of rDNA, which are species-informative.

The real-time PCR developed in this study is for diagnostic purposes only, and by no means intended for quantitative PCR (qPCR), because qPCR cannot be reliable in the case of pooled mosquitoes belonging to different species due to inter-specific variations in rDNA copy number [16] and body mass. However, the proportion of *An. stephensi* in a mosquito population, when present in extremely low density, can be obtained by the method employed for the estimation of infection rates in hematophagous insects by estimating minimum infection rate (MIR) or maximum likelihood procedure [17] based on the number of pools positive, methods frequently used for xenomonitoring.

LOD for the real-time PCR is considered an important criterion for assessing the sensitivity of real-time PCR, when the copy number of target nucleic acid is a limiting factor, for example, detecting pathogens in an organism. However, LOD is not a limiting factor for the *An. stephensi*-specific diagnostic real-time PCR, because rDNA is abundantly found in the organism due to its high copy number. It was observed in this study, that LOD cannot be a limiting factor even when the proportion of target mosquitoes is 1/500 in a pooled sample or tested on DNA isolated from a single leg (Ct values <24). Based on our observations, we suggest a cut-off value of 30 for real-time PCR, for the more reliable results. Ct values above this threshold can be suspected to be DNA contamination which should be verified through DNA sequencing using protocol described in **Supplementary Text S1**. In this study we observed false positivity for *An. stephensi* with late Ct values (>32) in two DNA samples (one each of the *Ae. aegypti* and *An. subpictus*) due to the contamination of DNA from *An. stephensi*.

The diagnostic PCRs in this study were designed to identify *An. stephensi* in large pools of samples. However, pooling of a large number of samples can accumulate potential PCR inhibitors. The heme compound in the blood [18] and eye pigment in the head of an insect [19] are reported potential inhibitors. Although we didn’t observe inhibitory effect of pooling in the real-time PCR assay, significant inhibitory effect was seen in a size-diagnostic PCR assay with a pool of 50 or more mosquitoes. In this study, we experienced an improvement in the intensity of the band in size-diagnostic PCR in such pools by diluting DNA.

Although we have successfully demonstrated identifying a single *An. stephensi* in pools of 500 mosquitoes, but it is recommended to use a pool of up to 100 mosquitoes that can be ground in a single microcentrifuge tube during DNA isolation, without the need for grinding by mortar and pestle. Grinding using a mortar and pestle may add chances of carryover contamination. For confirmation of *An. stephensi* in pooled samples through Sanger sequencing, as a precautionary measure, we suggest using internal primers (Stq-F and StD2-R, both of which are specific to *An. stephensi*) for sequence termination reactions. This is important to rule out sequencing of false-positive PCR products due to non-specific annealing with unknown nontarget species if any.

In conclusion, the molecular tools developed in this study can be used for the identification/confirmation of *An. stephensi*, individually or in a pool of mixed mosquito species enabling health authorities to detect early invasion of the species, especially in areas where it exists with low density.

## Availability of data

The dataset supporting the conclusions of this article is included within the article.

## Authors’ contribution

OPS designed PCR strategies, analysed data, wrote the first draft of the manuscript; GS, TK, KP and PKM performed laboratory experiments; PK, SM and NK contributed to the manuscript. All authors approved the final version of the manuscript.

## Ethics approval

Not required

## Acknowledgment

The following reagents were obtained through BEI Resources, NIAID, NIH: (1) *An. gambiae*, Strain G3, MRA-132K, contributed by Mark Q. Benedict, (2) *An. albimanus*, Strain STECLA, MRA-133K, contributed by Mark Q. Benedict, (3) *An. stephensi*, Strain STE2, MRA-134K, contributed by Mark Q. Benedict, (4) *An. freeborni*, Strain F-1, MRA-136K, contributed by Mark Q. Benedict, (5) *An. quadrimaculatus*, Strain ORLANDO, MRA-137K, contributed by Mark Q. Benedict, (6) *An. dirus*, Strain WRAIR2, MRA-700K, contributed by Mark Q. Benedict, (7) *An. farauti*, Strain FAR1, MRA-489K, contributed by Mark Q. Benedict, (8) *An. funestus*, Strain FUMOZ, MRA-1027K, contributed by Maureen Coetzee, (9) *An. merus*, Strain OPHANSI, MRA-803B, contributed by Rajendra Maharaj, and (10) *An. atroparvus*, Strain EBRO, MRA-493B, contributed by Carlos Aranda and Mark Q. Benedict. Authors are thankful to Kanwar Singh for mosquito collection, Priya Agrohi and Vijay Kumar for DNA sequencing and Abhinav Sinha for critical review of the manuscript.

