## Supplementary file for "Molecular diagnostics for early detection of invasion of malaria vector *Anopheles stephensi*"

**Supplementary information**

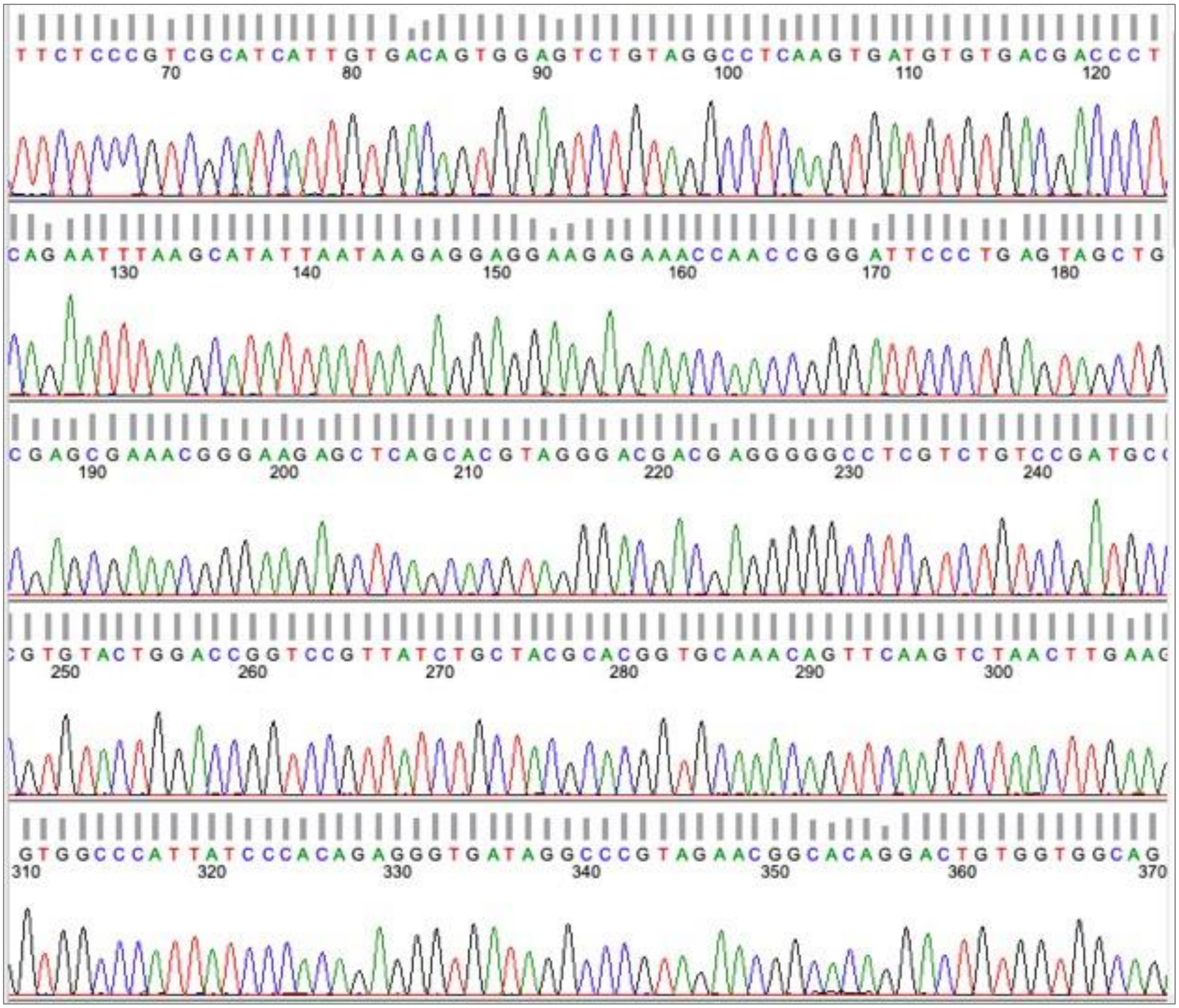

**Figure S1.** DNA sequence chromatogram (partial ITS2 and 28S) of *An. stephensi* showing the quality of sequence derived from DNA isolated from a pool of 500 mosquitoes containing single *An. stephensi* and rest *An. culicifacies.*

### Supplementary Text S1. DNA sequencing of mosquito samples detected false positive for *An. stephensi* in real-time PCR with late Ct value

Two pre-isolated DNA of mosquitoes, one each of *Aedes aegypti* and *An. subpictus* were detected false-positive as *An. stephensi* by real-time PCR with late Ct values (32.95 and 33.09, respectively). To check if false positivity is due to the contamination of DNA from *An. stephensi*, these samples were subjected to DNA sequencing following method described in this paper under section "DNA sequencing strategy for the confirmation of PCR-based identification of *An. stephensi* in pooled samples" with modification where number of PCR cycles were increased to 45. Both samples showed ~450 bp amplicon (**Figure S2**). The PCR products were sequenced from both directions of the strand. NCBI-blast search revealed 100% homology of all sequences with *An. stephensi*, confirming contamination of samples with DNA from *An. stephensi*.

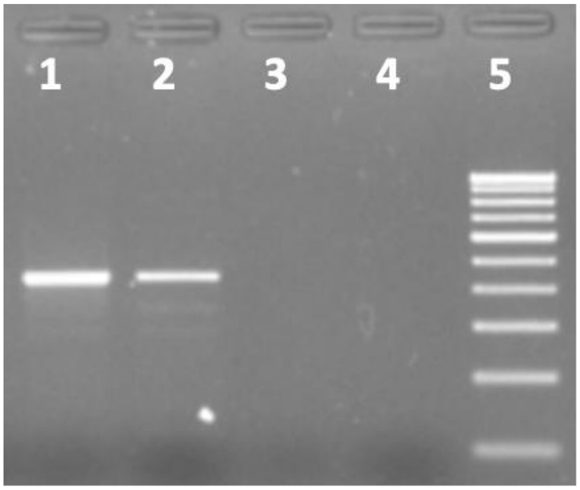


**Figure S2**. Gel photograph showing amplification of *An. stephensi*-specific PCR product in DNA isolated from *Ae. aegypti* (lane 1) and *An. subpictus* (lane 2) which were detected false positive for *An. stephensi* in real-time PCR with late Ct values. Lanes 3 and 4 are negative controls; Lane 5: 100 bp DNA ladder.
